# Strong Aversive Conditioning Triggers a Long-Lasting Generalized Aversion

**DOI:** 10.1101/2022.01.10.475691

**Authors:** Raul Ramos, Chi-Hong Wu, Gina G. Turrigiano

## Abstract

Generalization is an adaptive mnemonic process in which an animal can leverage past learning experiences to navigate future scenarios, but overgeneralization is a hallmark feature of anxiety disorders. Therefore, understanding the synaptic plasticity mechanisms that govern memory generalization and its persistence is an important goal. Here, we demonstrate that strong CTA conditioning results in a long-lasting generalized aversion that persists for at least two weeks. Using brain slice electrophysiology and activity-dependent labeling of the conditioning-active neuronal ensemble within the gustatory cortex, we find that strong CTA conditioning induces a long-lasting increase in synaptic strengths that occurs uniformly across superficial and deep layers of GC. Repeated exposure to salt, the generalized tastant, causes a rapid attenuation of the generalized aversion that correlates with a reversal of the CTA-induced increases in synaptic strength. Unlike the uniform strengthening that happens across layers, reversal of the generalized aversion results in a more pronounced depression of synaptic strengths in superficial layers. Finally, the generalized aversion and its reversal do not impact the acquisition and maintenance of the aversion to the conditioned tastant (saccharin). The strong correlation between the generalized aversion and synaptic strengthening, and the reversal of both in superficial layers by repeated salt exposure, strongly suggests that the synaptic changes in superficial layers contribute to the formation and reversal of the generalized aversion. In contrast, the persistence of synaptic strengthening in deep layers correlates with the persistence of CTA. Taken together, our data suggest that layer-specific synaptic plasticity mechanisms separately govern the persistence and generalization of CTA memory.

## 1 Introduction

The cellular and synaptic physiology underlying the formation, storage, and retrieval of memories has been extensively studied (Kandel et al., 2013; Kandel, Dudai, & Mayford, 2014; Josselyn & Tonegawa, 2020), but comparatively less research has focused on the synaptic plasticity mechanisms that govern the generalization of memories (Asok, Kandel, & Rayman, 2019). Generalization is an adaptive memory process by which an animal is able to extend its learning, garnered from past experiences, to future similar scenarios (Shepard, 1987; Asok, Kandel, & Rayman, 2019). In this way, generalization endows organisms with greater cognitive flexibility and makes them more savvy survivors. However, generalization can also be maladaptive, and overgeneralization of aversive memories has been implicated in post-traumatic stress disorder and anxiety disorders (Kheirbek et al., 2012; Mahan & Ressler, 2012; Dunsmoor & Paz, 2015). Thus, expanding our understanding of the synaptic plasticity mechanisms that shape memory generalization and its persistence is of great importance.

Conditioned Taste Aversion (CTA) learning is an ethologically relevant form of associative learning in which experience with a novel tastant (CS) is paired with LiCl induced gastric malaise (US) to produce a learned aversion (Bures, Bermúdez-Rattoni, & Yamamoto, 1998; Reilly & Schachtman, 2008). Unlike other associative learning paradigms that require multiple CS-US pairings, CTA results in robust and rapid learning following one conditioning trial (Bures, Bermúdez-Rattoni, & Yamamoto, 1998; Reilly & Schachtman, 2008). This feature of CTA learning makes it an ideal paradigm to explore how differences in conditioning strength might alter the temporal dynamics of memory generalization. The generalized aversion resulting from CTA conditioning is a well-documented phenomenon (Domjan, 1975; Parker & Revusky, 1982; Richardson, Williams, & Riccio, 1984; Smith & Theodore, 1984; Frank & Nowlis, 1989; Chotro & Alonso, 1999; Heyer et al., 2003; Baird, St. John, & Nguyen, 2005; Smith et al., 2012; Angulo, 2018; Wu, Ramos, et al., 2021), but it is unclear how long this generalization can persist, which factors impact its duration, and the cellular basis of this persistence.

Previously, we found that moderate conditioning (using a lower concentration of LiCl) induced a generalized aversion that reversed within 24 hours, while strong conditioning (using a higher concentration of LiCl) produced a generalized aversion that was still present at the 24 hours’ time point (Wu, Ramos, et al., 2021). Here, we explored this more persistent generalized aversion to determine how long it lasts, and the cellular basis of its persistence. We discovered that strong aversive conditioning results in a long-lasting generalized aversion that persisted for up to two-weeks and only reversed after experience with the generalized tastant. Next, we sought to characterize the cellular basis of its expression. We utilized the robust activity marking (RAM; Sørensen et al., 2016) system to label ensembles of neurons within gustatory cortex (GC) that were active during conditioning, and then performed *ex-vivo* whole-cell brain slice electrophysiology to measure excitatory postsynaptic strengths. Previous work has demonstrated the importance of the GC in CTA memory generalization (Kiefer & Braun, 1977; Kiefer & Braun, 1979; Wu, Ramos, et al., 2021). Our experiments revealed a potentiation of synaptic strengths occurring uniformly across the superficial and deep layers of GC in animals that had undergone strong CTA conditioning. Moreover, reversal of the long-lasting generalized aversion correlated with a reversal of the increases in synaptic strength. Unlike the CTA-induced potentiation that was observed uniformly across layers of GC, the exposure-induced reversal of synaptic potentiation occurred selectively in the superficial layers of GC. Together these findings illuminate the synaptic plasticity mechanisms that support a persistent long-lasting generalized aversion.

## 2 Materials and Methods

### 2.1 Experimental subject details

All experimental procedures were approved by Brandeis University Institutional Animal Care and Use Committee and followed the National Institute of Health guidelines for the Care and Use of Laboratory Animals. Young Long-Evans rats p28-p34 of both sexes were used in these experiments. Timed pregnant rats were obtained from Charles River Laboratories, and the progeny were maintained in the Foster Biomedical Research Labs at Brandeis University. After weaning at p21, littermates were individually housed in a humidity- and temperature-controlled environment and entrained to a 12-hour light-dark cycle (light phase from 7:00-7:00) with *ad libitum* access to food and water unless described otherwise. All subjects selected for electrophysiology experiments were age matched to animals selected for behavioral experiments.

### 2.2 Behavior

#### 2.2.1 Two-bottle paradigm

This CTA behavioral paradigm was conducted as previously described (Wu, Ramos, et al., 2021). Animals were transferred into individual home cages at p21 and habituated to two bottles with *ad libitum* access to water for three days. Next, the animals underwent water restriction for an additional three days, during which the access to water was limited to two hours. On the fourth day of restriction, rats were subjected to CTA conditioning. They received two bottles that contained the conditioned stimulus (CS, 10 mM saccharin), for thirty minutes, followed by an intraperitoneal injection of the unconditioned stimulus (US, LiCl). For moderate conditioning we used 0.15 M LiCl and for strong conditioning we used 0.30 M LiCl, dosed at 1ml/100g body weight (1% body weight). After conditioning, the water restriction schedule was continued until testing. For the two-bottle test, rats were given one bottle of tastant and one bottle of water, for thirty minutes. The location of the two bottles was counterbalanced across presentations to prevent any positional biases. The results were quantified using a tastant preference score (TPS):

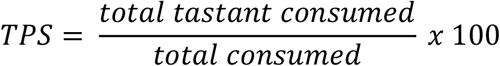

To test for a generalized aversion (Gen. test), salt (150 mM NaCl) was used as a tastant. To measure the reversal and attenuation of the generalized aversion, Gen. tests were conducted daily for three days. After Gen. testing was complete, rats were given a CTA test to ensure that they had indeed learned an aversion to the CS. All consumption was documented throughout the course of this behavior to ensure that daily fluid intake was stable. No rats were excluded.

### 2.3 RAM labeling of conditioning-active neurons

#### 2.3.1 Viral vector

RAM-dtTA-TRE-tdTomato was packaged in AAV serotype 9 by Duke Viral Vector Core.

#### 2.3.2 Virus surgery

Rats were anesthetized via intraperitoneal infusion of a cocktail containing ketamine (70 mg/kg), xylazine hydrochloride (3.5 mg/kg), and acepromazine maleate (0.7 mg/kg), and placed onto a stereotaxic apparatus. The stereotaxic apparatus was localized over a heating pad to maintain the animals body temperature. The skull was exposed and craniotomies were made above gustatory cortex (GC) using the following coordinates: anterior-posterior with reference to bregma: 1.0 mm; medial-lateral: ± 4.7 mm; dorsal-ventral with reference to the brain surface: 3.6 mm. RAM virus (400 nL) was unilaterally microinjected into GC through a glass micropipette connected to a micromanipulator (Narishige, MO-10), at a rate of approximately 200 nl/min. To allow adequate diffusion of virus particles, the pipet remained in place for additional 3 minutes after injection and was slowly withdrawn from the site. Injections were counterbalanced across hemispheres and both hemispheres are equally represented in the data.

### 2.3.3 Labeling of conditioning-active neurons

Customized chow containing low-dose doxycycline (40 ppm, ScottPharma) was added to the home cage one day before virus surgery, and rats were maintained on doxycycline until CTA conditioning. The doxycycline-containing chow was removed and replaced with regular chow one day before the conditioning trial to allow adequate RAM induction. Two hours after the conditioning trial, rats were placed back on the doxycycline diet to prevent further RAM activation. Acute brain slices were collected 3 days after conditioning.

### 2.4 Electrophysiology

#### 2.4.1 *Ex-vivo* acute brain-slice preparation

Brain slices were produced following our previously documented protocols (Miska et al., 2018; Cary & Turrigiano, 2021; Wu, Ramos, et al., 2021). Briefly, rats (p28-p34) were anesthetized with isoflurane, decapitated, and the brain was swiftly dissected out into ice cold carbogenated (95% O2, 5% CO2) standard ACSF (in mM: 126NaCl, 25 NaHCO3, 3 KCl, 2 CaCl2, 2 MgSO4, 1 NaH2PO4, 0.5 Na-Ascorbate, osmolarity adjusted to 310-315 mOsm with dextrose, pH 7.35). Coronal brain slices (300mm) containing GC were obtained from virus injected hemispheres using a vibratome (LeicaVT1000). The slices were immediately transferred to a warm (34°C) chamber filled with a continuously carbogenated ‘protective recovery’ choline-based solution (in mM: 110 Choline-Cl, 25 NaHCO3, 11.6 Na-Ascorbate, 7 MgCl2, 3.1 Na-Pyruvate, 2.5 KCl, 1.25NaH2PO4, and 0.5 CaCl2, osmolarity 310-315 mOsm, pH 7.35) for 10 minutes (Ting et al., 2014). After incubation in choline, the slices were transferred back to warm (34°C) carbogenated standard ACSF and incubated another 45 minutes. Brain slices were used for electrophysiology experiments between 1 - 7 hours post-slicing.

#### 2.4.2 Whole-cell patch clamp recording

Slices were visualized on an Olympus upright epifluorescence microscope using a 10x air (0.13 numerical aperture) and 40x water-immersion objective (0.8 numerical aperture) with infrared-differential interference contrast optics and an infrared CCD camera. Gustatory cortex was identified in acute slices using the shape and morphology of the corpus callosum, piriform cortex and the lateral ventricle as a reference. The borders of GC were determined by comparing the aforementioned landmarks in slice to the Paxinos and Watson rat brain atlas. Pyramidal neurons from superficial and deep layers across the dorsal-ventral axis of GC were visually targeted and identified by the presence of an apical dendrite and teardrop shaped soma. In experiments involving the expression of a viral construct, such as RAM, fluorophore expression was used to visually target pyramidal neurons. Virus expression was consistent across layers (Wu, Ramos, et al., 2021). Pyramidal morphology was confirmed by post hoc reconstruction of biocytin fills. Borosilicate glass recording pipettes were pulled using a Sutter P-97 micropipette puller, with acceptable tip resistances ranging from 3 to 6 Mn. All recordings were performed on submerged slices, continuously perfused with carbogenated 35°C recording solution. Data were low-pass filtered at 10 kHz and acquired at 10 kHz with Axopatch 700B amplifiers and CV-7B headstages (Molecular Devices, Sunnyvale CA). Data were acquired using WaveSurfer v0.953 (Janelia Research Campus), and all post hoc data analysis was performed using in-house scripts written in MATLAB (Mathworks, Natick MA).

#### 2.4.3 mEPSC recordings

Cs+ Methanesulfonate-based internal recording solution was used as previously reported (Miska et al., 2018; Wu, Ramos, et al., 2021). This Cs+ internal contained (in mM) 115 Cs-Methanesulfonate, 10 HEPES, 10 BAPTA•4Cs, 5.37 Biocytin, 2 QX314 Cl, 1.5MgCl2, 1 EGTA, 10 Na2-Phosphocreatine, 4 ATP-Mg, and 0.3 GTP-Na, with sucrose added to bring osmolarity to 295 mOsm, and CsOH added to bring pH to 7.35. For these recordings, pyramidal neurons were voltage clamped to -70 mV in standard ACSF containing a drug cocktail of TTX (0.2mM), APV (50mM), PTX (25mM). Traces of 10 s were acquired over a period of ∼10-15 minutes allowing for the cell to fill for later morphological verification. Neurons were excluded from analysis if Rs > 25 Mn or if was Rin > 2a above the mean.

#### 2.4.4 mEPSC analysis

To reliably detect mEPSC events and limit selection bias, we used in-house software that employs a semi-automated template-based detection method (Miska et al., 2018; Cary & Turrigiano, 2021).

Event inclusion criteria included amplitudes greater than 5 pA and rise times less than 3 ms. The resulting events detected by our software were visually assessed posthoc and a subset of artefactual events were excluded. The experimenter was blinded to experimental condition and treatment until after the analysis was complete.

#### 2.4.5 Recording from the conditioning-active ensemble

Slices were collected exactly 3-days post-conditioning using the methods described above. mEPSCs were recorded using the method described above. Fluorescent RAM^+^ (tdTomato^+^) cells were targeted in both superficial and deep layers, where expression was equally robust (Wu, Ramos, et al., 2021). The expression, or lack of, tdTomato was confirmed post hoc through immunostaining of the cells using antibodies described in the immunohistochemistry section.

### 2.5 Immunohistochemistry

#### 2.5.1 Immunostaining of biocytin-filled cells

After recording, slices were incubated in cold 4% PFA for two days to fix the tissue. Following fixation, slices were washed six times with PBS, preincubated with blocking buffer (5% goat serum, 3%BSA, 0.3% Triton X-100 in PBS) for three hours at room temperature, and then incubated with primary antibodies diluted in diluent buffer (5% goat serum, 3%BSA in PBS) at 4°C overnight. RAM expression was verified using rabbit polyclonal anti-RFP (1:1000; Rockland 600-401-379). Slices were counterstained with mouse monoclonal anti-NeuN (1:500; Millipore MAB-377). Following incubation with primary antibodies, slices were washed six times with PBS and then incubated at 4°C overnight with secondary antibodies diluted in diluent buffer. The secondary antibodies used for these experiments were goat polyclonal anti-rabbit Alexa Fluor 568 (1:500; Thermo-Fisher, A-11036) and goat polyclonal anti-mouse Alexa Fluor 647 (1:500; Thermo-Fisher, A-21236). Biocytin fills were recovered by staining with streptavidin Alexa Fluor 488 (1:350; Thermo-Fisher, S-11223). After incubation with secondary antibodies, slices were washed and mounted for imaging. Images were acquired using the Leica SP5Laser Scanning Confocal Microscope.

### 2.6 Quantification and statistics

For all experiments including behavior and electrophysiology, individual experimental distributions were tested for normality using the Anderson-Darling test. All experimental conditions passed the normality test. A one sample, or two sample t test, or one-way ANOVA were used where appropriate. Significant ANOVA tests were followed by Tukey-Kramer post hoc comparisons. Results of all statistical tests can be found in the figure legends. For behavior experiments n = number of animals, while for electrophysiology experiments n = number of cells. Electrophysiology data were collected from at least 4 animals for each condition. Scatterplots were generated using publicly available MATLAB code (Ramirez, 2020).

## 3 Results

### 3.1 Strong aversive conditioning triggers a long-lasting generalized aversion

We previously demonstrated that strong CTA conditioning results in a generalized aversion that could be observed at 24 hours post-conditioning (Wu, Ramos, et al., 2021), but it was unclear whether this generalized aversion persists beyond this time point. Therefore, we first sought to delineate how long the generalized aversion induced by strong CTA persists. We took animals through strong CTA conditioning (strong CTA, 0.30M LiCl) or, as a control, a more moderate conditioning regime (moderate CTA, 0.15M LiCl) which only induces a transient generalized aversion that subsides within the first 24 hours post-conditioning (Wu, Ramos, et al., 2021). We then tested for a generalized aversion 3 days post-conditioning (**Figure 1A**). We found that strong CTA training resulted in a significant generalized aversion to salt at 3 days post-conditioning compared to moderate CTA controls (**Figure 1B**). Repeated presentations of salt in the strong CTA group rapidly reversed the generalized aversion, attenuating after just one presentation (**Figure 1C**). By contrast, in the moderate CTA group which does not exhibit a long-lasting generalized aversion, repeated presentation of salt caused no significant change in tastant preference score (**Figure 1D**). The expression of the generalized aversion, as well as its reversal, did not impact the specific taste aversion to the conditioned tastant, as animals still showed a strong aversion to saccharin after generalization tests were completed (**Figure 1E**). These findings are consistent with our previous results at the 24-hour time point for both experimental conditions (Wu, Ramos, et al., 2021).

**Figure 1.**
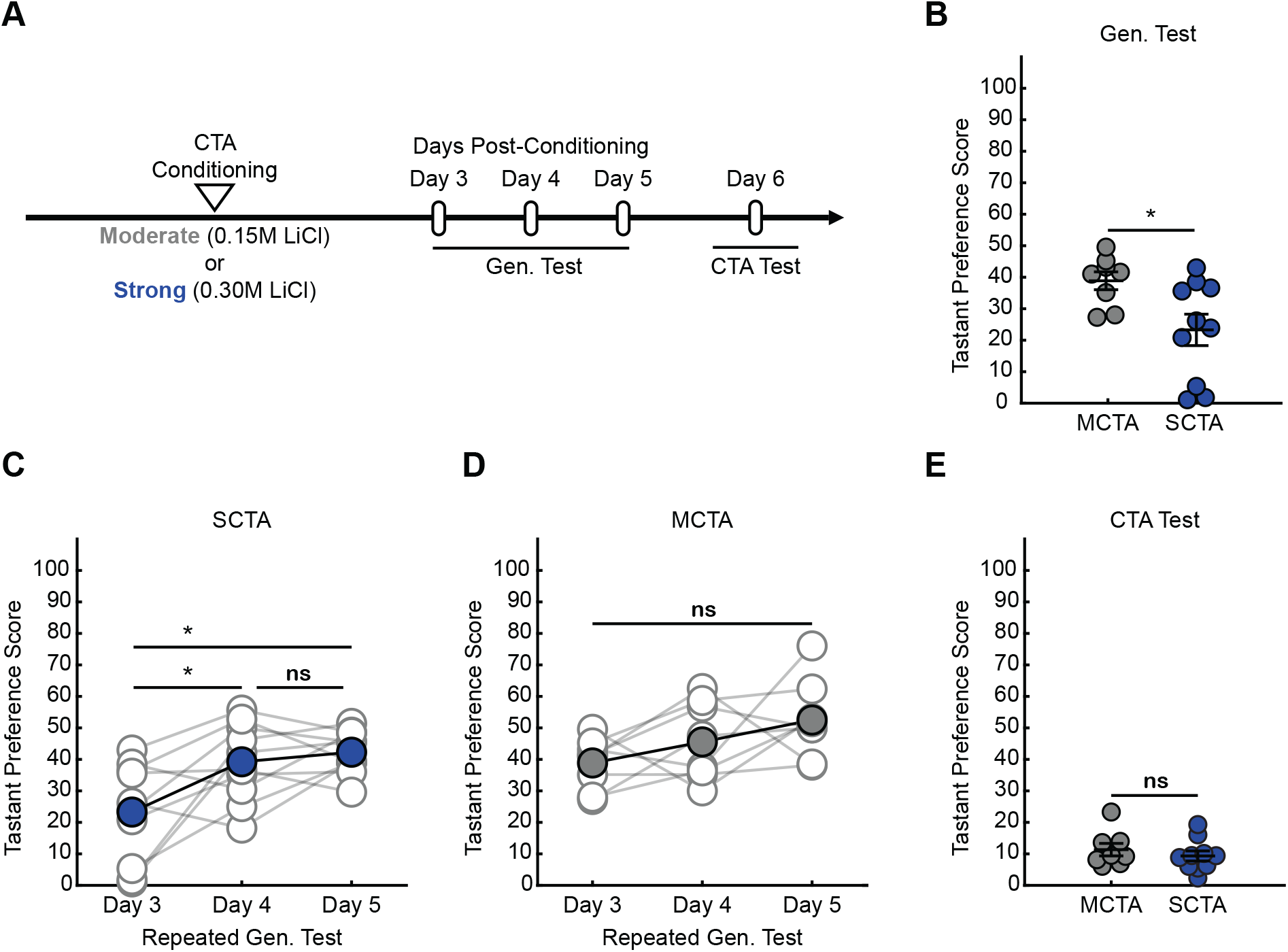
Strong aversive conditioning triggers a long-lasting generalized aversion. (**A**) Two-bottle CTA learning paradigm for testing the persistence of the generalized aversion. (**B**) Gen. test; preference score for salt tested 3 days post-conditioning (two-sample t test, p = 0.0220). (**C**) SCTA repeated Gen. test; preference for salt tested at 3-, 4-, and 5-days post-conditioning (One-way ANOVA, p = 0.0036; Tukey-Kramer post-hoc tests, day 3 vs day 4, p = 0.0184; day3 vs day 5, p = 0.0047; day 4 vs day 5, p = 0.8430). (**D**) MCTA repeated Gen. test; preference for salt tested at 3-, 4-, and 5-days post-conditioning (One-way ANOVA, p = 0.0716). (**E**) CTA test; preference score for saccharin of 3-day experimental conditions (two-sample t test, p = 0.4387).

Subsequent experiments revealed the long-lasting nature of the generalized aversion. We found that we could observe a generalized aversion resulting from strong CTA conditioning at 7-, 10-, and 14-days post-conditioning (**Figure 2A & 2B**). In all instances, the generalized aversion reversed after one trial with the generalized tastant without impacting the specific taste aversion to saccharine (**Figure 2B & 2C**). Together, these results demonstrate that increasing the strength of conditioning with a higher concentration of LiCl triggers a long-lasting generalized aversion that persists up to two weeks post-conditioning and only reverses after experience with the generalized tastant.

**Figure 2.**
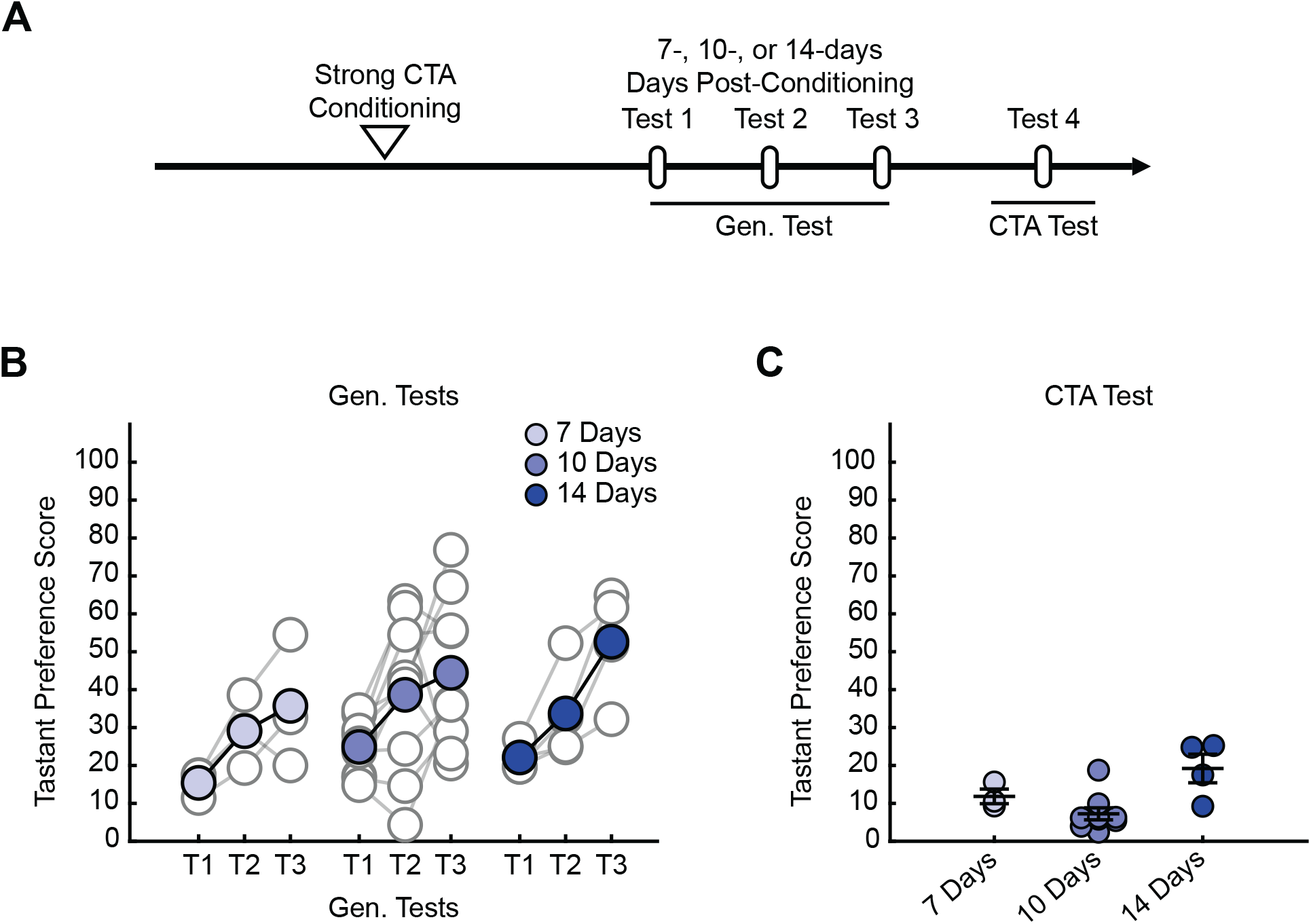
Strong CTA-induced generalized aversion can be observed up to two-weeks post-conditioning (**A**) Two-bottle CTA learning paradigm for testing the persistence of the generalized aversion at 7-, 10-, and 14-days post-conditioning. (**B**) Repeated Gen. test; preference score for NaCl tested 7-, 10-, and 14-days experimental conditions. Each test is separated by a 24-hour interval. (**C**) CTA test; preference score for saccharin of long-duration experimental conditions.

### 3.2 Strong aversive conditioning drives a long-lasting increase in postsynaptic strength

We next endeavored to find an electrophysiological correlate for this long-lasting generalized aversion within the gustatory cortex. Previously, we found that in the conditioning-active GC neurons, moderate CTA conditioning produced a transient increase in postsynaptic strengths that was homeostatically scaled down, while strong CTA conditioning resulted in a more sustained increase of postsynaptic strengths (Wu, Ramos, et al., 2021). Notably, the presence or absence of this increase in postsynaptic strengths correlated with the presence or absence of the generalized aversion. We thus hypothesized that a long-lasting increase in postsynaptic strengths could support the long-lasting generalized aversion induced by strong CTA conditioning. To test this, we virally expressed the robust activity marking (RAM) system in GC to label the conditioning-activated neuronal ensemble through the activity dependent expression of tdTomato (Sørensen et al., 2016; Wu, Ramos, et al., 2021). RAM labeling is inhibited by doxycycline (Dox); by removing Dox we could restrict labeling to a small window during which animals underwent either strong or moderate CTA conditioning.

Congruent with the behavioral measurements in **Figure 1**, brain slices were prepared 3 days after CTA induction for electrophysiological recording (**Figure 3A**). We then targeted either RAM^+^ (tdTomato^+^) or nearby RAM^-^ (tdTomato^-^) pyramidal neurons for whole-cell electrophysiological recordings (**Figure 3B**) and measured miniature excitatory postsynaptic currents (mEPSCs) to quantify postsynaptic strength (**Figure 3C**). We found that strong CTA conditioning produced a significant increase in mEPSC amplitude in RAM^+^ neurons; more surprisingly, a similar increase was observed in RAM^-^ neurons when compared to neurons recorded from moderate CTA animals (**Figure 3D & 3E**). Other studies have similarly documented changes in synaptic strength in GC neurons using whole-cell physiology following CTA learning without an activity-dependent labeling scheme, suggesting that synaptic plasticity occurs across a large population of cells in GC (Haley et al., 2020). These findings reveal that strong CTA conditioning produces a long-lasting increase in postsynaptic strengths that parallels the long-lasting generalized aversion and suggest a possible GC-wide induction of synaptic plasticity. Because neurons can be RAM^-^ either because they did not express the virus, or because they did but were not active during conditioning, for the remainder of our experiments we focused solely on RAM^+^ neurons to ensure we recorded from a more homogenous population of cells.

**Figure 3.**
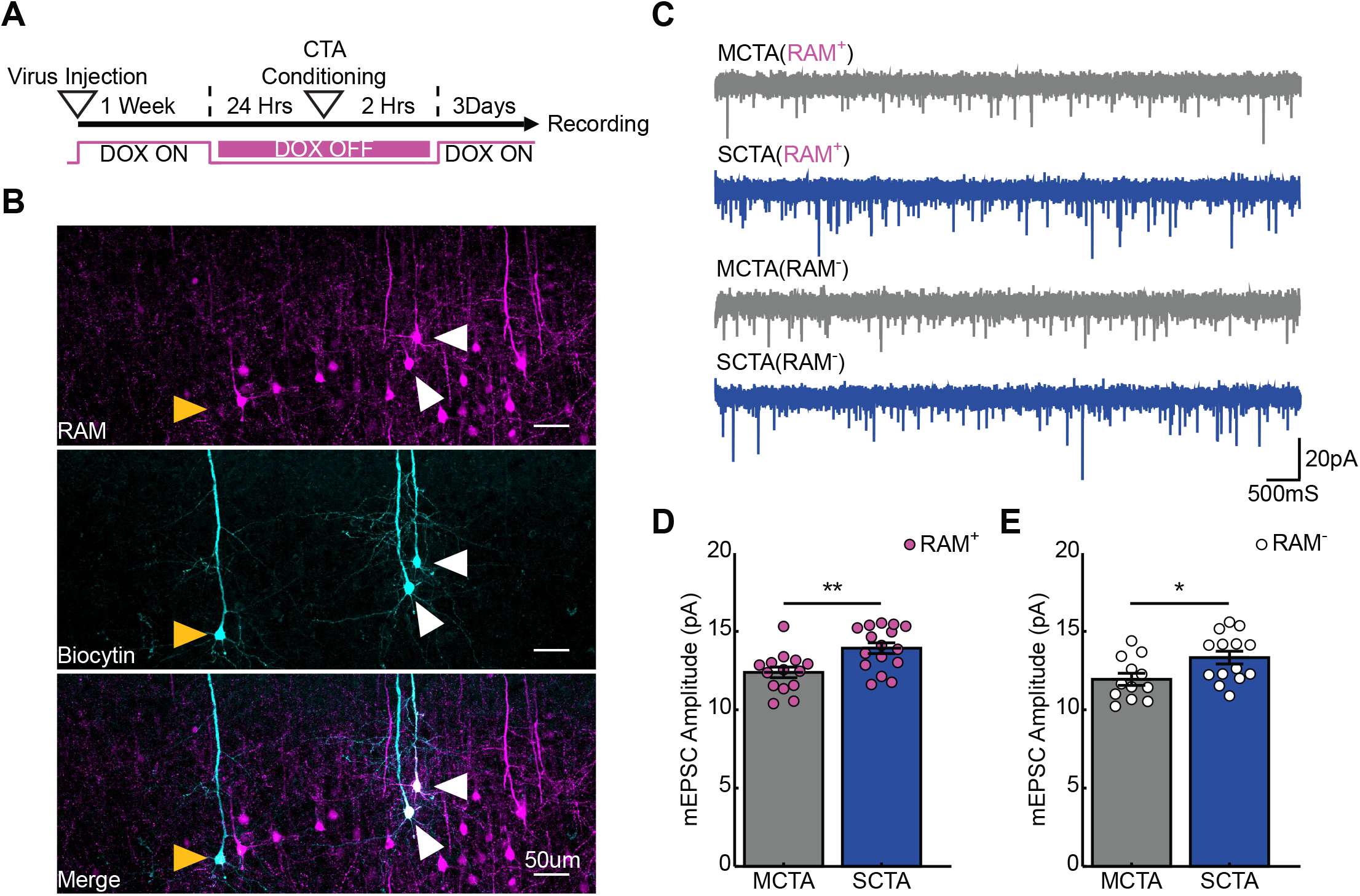
Strong aversive conditioning drives a long-lasting increase in postsynaptic strength. (**A**) Experimental design. (**B**) Biocytin fills (cyan) of RAM^+^ (magenta; white arrows) and RAM^-^ (yellow arrows) pyramidal cells in gustatory cortex. (**C**) Representative mEPSC recordings. (**D**) Cell-average mEPSC amplitudes of RAM^+^ cells 3 days post-conditioning (two-sample t test, p = 0.0035). (**E**) Cell-average mEPSC amplitudes of RAM^-^ cells 3 days post-conditioning (two-sample t test, p = 0.0224).

### 3.3 Experience with salt rapidly reverses increases in postsynaptic strength

If the increase in postsynaptic strength contributes to the expression of the long-lasting generalized aversion, then reversal of the generalized aversion by salt exposure should also reverse the increases in mEPSC amplitude in conditioning-active GC neurons. To test this hypothesis, animals were exposed to salt and then 2- or 24-hours following salt exposure, slices were made for electrophysiological recordings (**Figure 4A**). As in the previous experiments, at the time of recording, 3 days had elapsed post-conditioning. These experiments revealed that, when compared to the average baseline amplitude following strong CTA conditioning in **Figure 3** (green dashed line), experience with the generalized tastant (salt) rapidly decreased mEPSC amplitude. This depression was evident as early as 2 hours following salt presentation and was greater in magnitude after 24 hours (**Figure 4B**). In contrast, when compared to the average baseline amplitude following moderate CTA conditioning in **Figure 3** (green dashed line), animals in the moderate CTA group showed no changes at 2- or 24-hours post-salt exposure (**Figure 4C**). These findings reveal a strong correlation between postsynaptic strength and memory generalization within the conditioning-active ensemble. Furthermore, they suggest an important contribution from forms of synaptic depression that operate over a time scale of hours.

**Figure 4.**
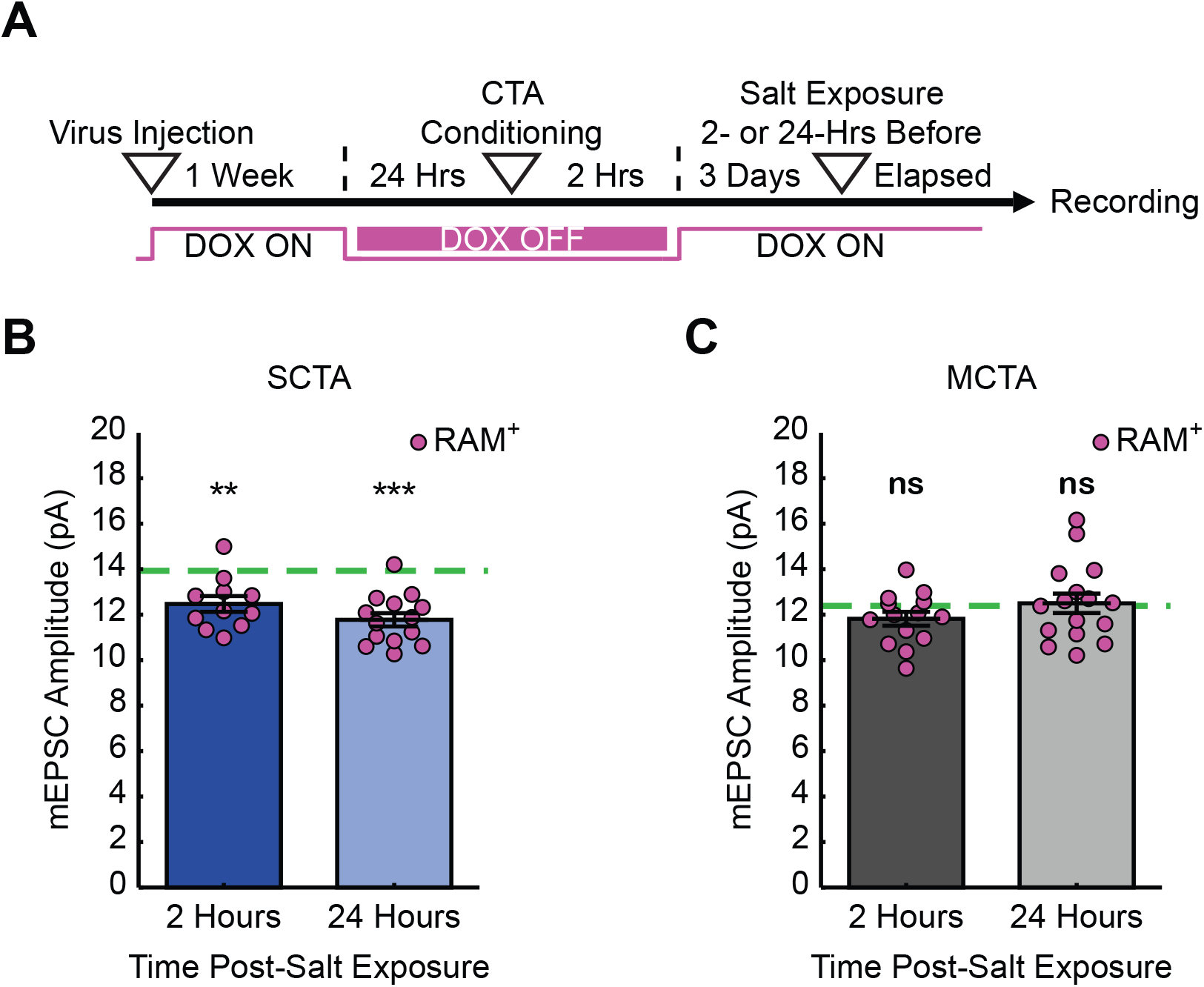
Experience with salt rapidly reverses increases in postsynaptic strength. (**A**) Experimental design. (**B**) SCTA cell-average mEPSC amplitudes of RAM^+^ cells 2- and 24-hours post-salt exposure. Green dashed line represents average mEPSC amplitude from RAM^+^ cells in SCTA no-salt exposure conditions (**Figure 3D**; one-sample t test against hypothesized mean, 2 hours, p = 0.0018, 24 hours, p = 5.5750e-06). (**C**) MCTA cell-average mEPSC amplitudes of RAM^+^ cells 2- and 24-hours post-salt exposure. Green dashed line represents average mEPSC amplitude from RAM^+^ cells in MCTA no-salt exposure conditions (**Figure 3D**; one-sample t test against hypothesized mean, 2 hours, p = 0.0857, 24 hours, p = 0.7945).

### 3.4 CTA-induced increase in postsynaptic strength occurs uniformly across superficial and deep layers of GC

It is known that superficial (layers II/III) and deep (layers V/VI) layers of the gustatory cortex are innervated differentially by structures important for CTA learning (Allen et al., 1991; Nakashima et al., 2000; Maffei, Haley, & Fontanini, 2012; Haley, Fontanini, & Maffei, 2016). This raised the question of whether synaptic strengthening following CTA learning is uniformly expressed across the different layers of gustatory cortex. The recordings presented in **Figure 3** were performed across superficial and deep layers of the GC (**Figure 5A**). When we sorted these data based on layers, we found that strong CTA conditioning indeed increased mEPSC amplitude (compared to moderate CTA controls) in both superficial and deep layers (**Figure 5B**). Thus, while these different layers are distinctly innervated, strong CTA induces widespread and uniform changes in synaptic plasticity across the cortical layers of GC.

**Figure 5.**
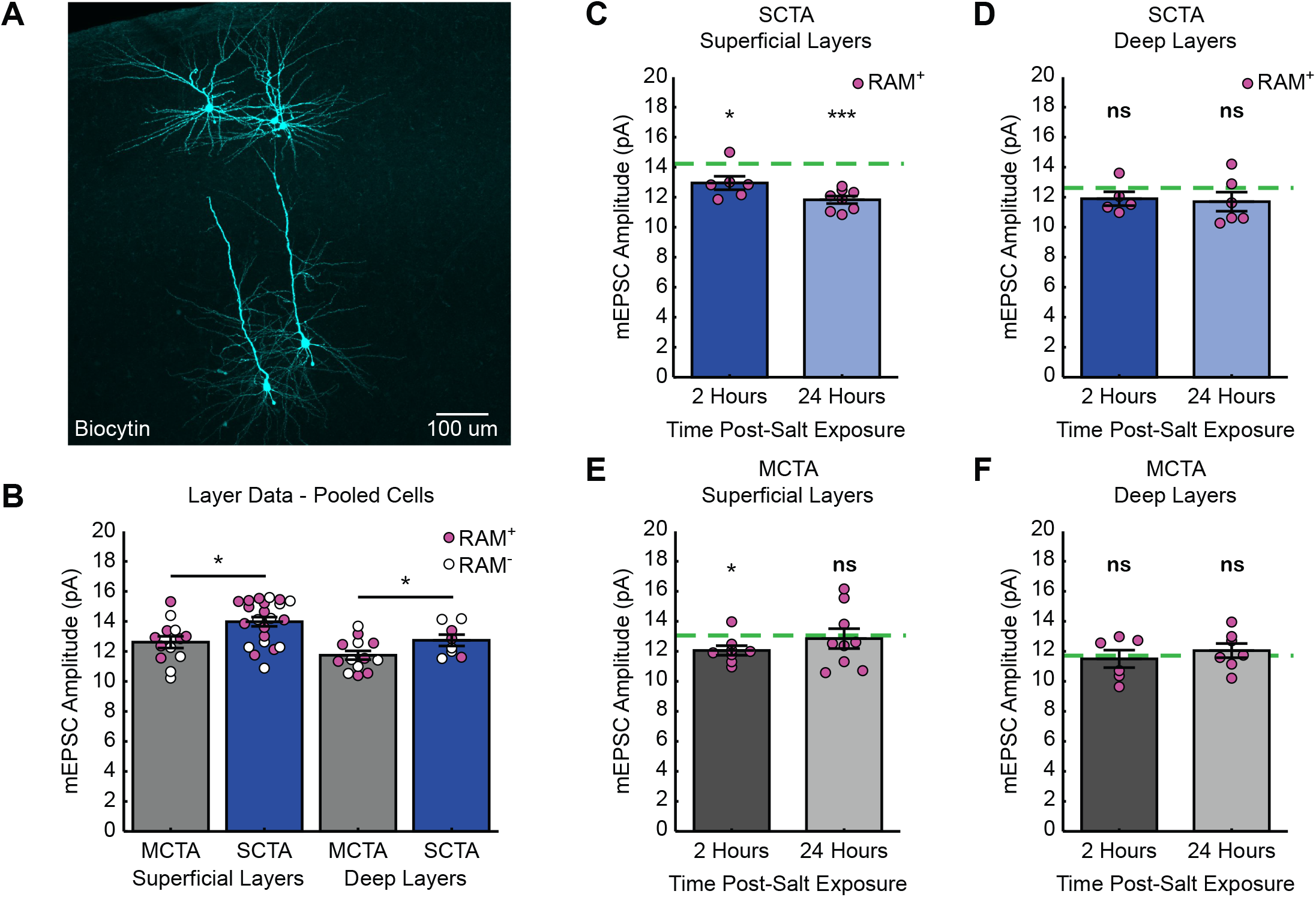
Distinct layer processes govern the induction and reversal of generalization. (**A**) Biocytin fills (cyan) of cells recorded from the superficial and deep layers of gustatory cortex. (**B**) Cell-average mEPSC amplitudes of pooled RAM^+^ and RAM^-^ cells 3 days post-conditioning sorted by superficial and deep layers (superficial layers, two-sample t test, p = 0.0104; deep layers, two-sample t test, p = 0.0498). (**C**) SCTA cell-average mEPSC amplitudes of RAM^+^ cells from superficial layers 2- and 24-hours post-salt exposure. Green dashed line represents average mEPSC amplitude from RAM^+^ cells from superficial layers in SCTA no-salt exposure conditions (**Figure 5B**; one-sample t test against hypothesized mean, 2 hours, p = 0.0355, 24 hours, p = 2.7205e-05). (**D**) SCTA cell-average mEPSC amplitudes of RAM^+^ cells from deep layers 2- and 24-hours post-salt exposure. Green dashed line represents average mEPSC amplitude from RAM^+^ cells from deep layers in SCTA no-salt exposure conditions (**Figure 5B**; one-sample t test against hypothesized mean, 2 hours, p = 0.1929, 24 hours, p = 0.2087). (**E**) MCTA cell-average mEPSC amplitudes of RAM^+^ cells from superficial layers 2- and 24-hours post-salt exposure. Green dashed line represents average mEPSC amplitude from RAM^+^ cells from superficial layers in MCTA no-salt exposure conditions (**Figure 5B**; one-sample t test against hypothesized mean, 2 hours, p = 0.0166, 24 hours, p = 0.7606). (**F**) MCTA cell-average mEPSC amplitudes of RAM^+^ cells from deep layers 2- and 24-hours post-salt exposure. Green dashed line represents average mEPSC amplitude from RAM^+^ cells from deep layers in MCTA no-salt exposure conditions (**Figure 5B**; one-sample t test against hypothesized mean, 2 hours, p = 0.7262, 24 hours, p = 0.5056).

### 3.5 Reversal is more pronounced in superficial layers of GC

We next asked if the reversal of postsynaptic strengths after salt exposure also occurred uniformly across superficial and deep layers. We sorted the dataset in **Figure 4** by layers and normalized each group after salt exposure to the corresponding average amplitude without salt exposure (green dashed line). This revealed that neurons in the superficial layers of gustatory cortex exhibited a reversal of mEPSC amplitude after salt exposure (**Figure 5C**). In contrast, although strong CTA induced an increase in mEPSC amplitude across both superficial and deep layers, the reduction in mEPSC amplitude after salt exposure was less pronounced in the deep layers and did not reach statistical significance (**Figure 5D**). In animals that underwent moderate CTA conditioning, there were little-to-no changes in mEPSC amplitude. In the superficial layers, there was a reduction at 2 hours that was gone at 24 hours, and in the deep layers there was no change at either time point (**Figure 5E & 5F**). Together these results indicate that reversal of postsynaptic strengths induced by salt exposure is a layer-specific process that primarily occurs within the superficial layers of GC.

## 4 Discussion

Understanding the cellular basis of memory generalization can yield important insights into the maladaptive, persistent overgeneralization that is characteristic of anxiety disorders. Here we took advantage of CTA learning’s one trial nature to study a long-lasting generalized aversion resulting from a single strong conditioning event. In doing so, we uncovered changes in postsynaptic strength occurring in a layer specific manner within the gustatory cortex that correlated with many of our behavioral observations. Our experiments revealed that postsynaptic strength in the superficial layers of GC seems to increase, then decrease, with the presence and then reversal of the long-lasting generalized aversion, respectively. Interestingly, postsynaptic strength in the deep layers of GC also increases after CTA learning, but does not significantly change as the generalized aversion reverses, but instead persists much like the specific taste aversion to saccharin. These data suggest that rather than being functionally homogenous, neurons within the conditioning active ensemble in GC subserve different memory functions depending on what layer they are found in. These results are in line with the functionally distinct engram ensembles found in the hippocampus that govern the balance between memory generalization and specificity (Sun et al., 2020). Future experiments are needed to identify exactly what form of synaptic plasticity is responsible for the CTA-induced strengthening, the synaptic depression induced by exposure to the generalized tastant, and whether these forms of synaptic plasticity are causally involved in this behavior.

In these experiments we leveraged of our ability to manipulate the strength of the learned aversion by changing the concentration of LiCl and taking animals through a strong aversive conditioning scheme (0.30M LiCl, 1% b.w.). This concentration and dosage of LiCl has been previously demonstrated to result in a maximum aversion to the CS (Nachman & Ashe, 1973). Using this approach, we found that after a single conditioning trial, animals can form a long-lasting generalized aversion that persists up to two weeks post-CTA training. This persistence, to our knowledge, is longer than any reported thus far in the CTA literature on generalization (Domjan, 1975; Parker & Revusky, 1982; Richardson, Williams, & Riccio, 1984; Smith & Theodore, 1984; Frank & Nowlis, 1989; Chotro & Alonso, 1999; Heyer et al., 2003; Baird, St. John, & Nguyen, 2005; Smith et al., 2012; Angulo, 2018). The ethological significance of this behavior and the mechanism by which this generalization is occurring remain to be worked out. One possibility is that strong aversive conditioning results in a state of long-lasting enhanced caution towards novel stimuli to protect the animal from further toxicosis. This may reflect toxicosis induced enhanced neophobia, where CTA produces enhancement to an already decreased intake of unfamiliar, novel stimuli (Domjan, 1975; Reilly & Schachtman, 2008; Lin et al., 2012). In the context of associative learning, generalization is thought to occur more easily across stimuli that share “elements” with each other (McLaren & Mackintosh, 2002). But what these elementary features are with respect to tastants and CTA is unknown. In addition to basic taste modality (sweet, sour, salty, etc..) the GC has been shown to encode other taste features such as identity, palatability, and novelty/familiarity, all of which are elements to potentially evaluate taste stimuli by (Gallo, Roldan, & Bures, 1992; Lin et al., 2012; Flores et al., 2018; Mukherjee, Wachutka, & Katz, 2019). In our study, what elementary features are shared by salt, the generalized tastant, and saccharin, the conditioned stimulus, remain to be identified. The results of future experiments will further our understanding regarding whether this is truly stimulus generalization between saccharin and salt, or an avoidance like behavior similar to enhanced neophobia.

Our electrophysiology experiments found that the long-lasting generalized aversion resulting from strong CTA correlated with a similarly persistent increase in postsynaptic strengths in both RAM^+^ and RAM^-^ cells. There are several possible interpretations regarding why RAM^-^ cells, many of which are presumably not activated during conditioning, also exhibit an increase in postsynaptic strengths post-conditioning. One possibility is that these cells could be infected with RAM virus, but expression of tdTomato is below the threshold required for detection during live-cell recording. This is unlikely because in all our experiments, cells were recovered post-hoc using biocytin fills and stained against tdTomato, which would amplify the signal and allow us to detect even very low levels of RAM expression. Another possibility is that these cells are not transfected with the virally expressed RAM system and are activated by conditioning. We cannot exclude this possibility using our current methodology. However, it is unlikely that all RAM^-^ neurons fall into this category, given that our recordings were performed on neighboring RAM^+^ and RAM^-^ neurons within cortical areas showing high tdTomato expression. The last possibility is that these are cells infected with RAM virus, but not activated during conditioning (true RAM^-^). It is possible that RAM^-^ cells are later recruited during memory consolidation, thereby manifesting changes in synaptic strength. CTA conditioning is known to induce dynamic changes in firing rate within GC, which may suggest that different neuronal populations may undergo changes in synaptic plasticity on distinct time scales post-conditioning and this temporal activation facilitates the formation of an aversive memory (Moran & Katz, 2014; Arieli, Younis, & Moran, 2021). Additionally, previous studies have documented changes in synaptic strength in GC neurons following CTA training without an activity dependent labeling scheme, suggesting that synaptic plasticity occurs across a large population of cells in GC (Haley et al., 2020). Taken together, these findings hint at the possibility of GC-wide induction of synaptic plasticity following CTA conditioning. The strong correlation between generalized aversion and this strengthening, including the ability to reverse both by salt exposure, suggests that this strengthening could be causally involved in the expression of the generalized aversion, but more experiments are needed to establish causality.

Our previous experiments revealed that synaptic downscaling of postsynaptic strengths was important for reversing the transient generalized aversion resulting from moderate CTA training, and establishing CTA specificity (Wu, Ramos, et al., 2021). Whether a similar mechanism is involved in the experience dependent reversal of the long-lasting generalized aversion is unknown. The experiments presented in **Figure 4** demonstrate that as soon as 2 hours following salt exposure, mEPSC amplitudes are significantly decreased. The decrease in mEPSC amplitude is more pronounced 24 hours after salt exposure, suggesting that this depotentiation of postsynaptic strengths is initiated rapidly and continues to unfold slowly over the course of 24 hours. No changes were observed following salt exposure in animals that underwent moderate CTA conditioning. These data reveal a strong correlation between the experience dependent reversal of the long-lasting generalized aversion and reversal of the increases in postsynaptic strength following salt exposure. Several forms of synaptic plasticity could potentially drive this decrease in postsynaptic strength. First, previous research has demonstrated that LTD is involved in the extinction of CTA (Li et al., 2016), and salt exposure may induce LTD in a similar fashion during reversal of the generalized aversion.

Alternatively, strong CTA conditioning may induce cellular mechanisms that over-power homeostatic synaptic scaling and prevent the transition from generalized to specific taste aversion. In this scenario, downscaling becomes “gated” by future tastant experiences. Recent research on the behavioral-state gating of homeostatic plasticity supports this possibility (Hengen et al., 2016; Torrado Pacheco et al., 2021). Future experiments that use antagonists targeting specific forms of synaptic plasticity will clarify the mechanisms that drive the reversal of postsynaptic strengths after salt exposure and could reveal important insights into how we can reverse or weaken more persistent forms of memory generalization.

GC consists of superficial and deep layers that receive distinct projections from sub-cortical nuclei important for CTA learning. For example, inputs from the gustatory thalamus have been shown to be uniformly diffused across the GC, with a slight bias for the granular layer (layer IV) and layer V (Allen et al., 1991; Nakashima et al., 2000; Maffei, Haley, & Fontanini, 2012). In contrast, the amygdala projects to both superficial and deep layers of GC, but the magnitude of these inputs is larger in the superficial layers (Haley, Fontanini, & Maffei, 2016). Compared to other sensory cortices, such as visual cortex, little work has been done to characterize the synaptic properties of distinct inputs into distinct layers of GC (Maffei, Haley, & Fontanini, 2012). Even less is known about how these different inputs are modulated by CTA learning and whether there exist layer specific motifs. Our finding that strong CTA conditioning produces increases in postsynaptic strengths in RAM^+^ and RAM^-^ neurons in both superficial and deep layers further supports the notion that CTA results in a GC-wide induction of synaptic plasticity. During reversal, the reduction in postsynaptic strength is more pronounced in superficial layers of GC. In comparison, little to no change was observed across layers in the moderate CTA animals following salt exposure. Altogether, these data suggest that while CTA learning induces region-wide changes in postsynaptic strengths, a layer specific plasticity mechanism may be responsible for the establishment and reversal of the generalized aversion. Given that superficial layers receive inputs from multiple subcortical nuclei, it will require further experiments to isolate specific populations of synaptic inputs in which the generalized tastant-induced changes in synaptic plasticity occur.

Experiments on short- and long-term memory have previously revealed that these different types of memory, operating over distinct time scales, produce short- and long-term changes in synaptic efficacy, respectively (Kandel et al., 2013; Kandel, Dudai, & Mayford, 2014). Whether similar mechanisms are at work in the context of memory generalization remained unknown. Our previous experiments using moderate CTA conditioning revealed that a transient generalized aversion was correlated with transient changes in synaptic strength, analogous to the short-term changes observed during short-term memory (Wu, Ramos, et al., 2021). Here, we found that the long-lasting generalized aversion resulting from strong CTA correlated with a similarly long-lasting increase in postsynaptic strengths, analogous to changes seen in long-term memory formation (Kandel, 2012; Kandel, Dudai, & Mayford, 2014; Takeuchi, Duszkiewicz, & Morris, 2014). Finally, exposure to salt rapidly reversed the generalized aversion without impacting the retrieval of CTA memory at later time points, suggesting that the encoding of CTA and the generalized aversion either use distinct mechanisms or occur in distinct populations of cells. Our data provides evidence for the latter and suggests that the different layers of GC play functionally distinct roles in CTA memory. These findings suggest an interesting hypothesis: that changes within the conditioning-active ensemble are important for the initial acquisition of CTA memory, but after acquisition, the CTA conditioning-active neuronal ensemble functionally diverges, and cells in the superficial layers instead work to support the generalized aversion while cells in the deep layers are dedicated to the specific taste aversion learned against the CS. This hypothesis is supported by evidence from studies in the hippocampus where it was found that within an engram ensemble there exists functionally heterogeneous populations of neurons that work to govern the balance of memory generalization and specificity (Sun et al., 2020). Furthermore, these functionally distinct ensembles are genetically defined based on their immediate early gene expression profile, either *Npas4* or *Fos*, both of which are transcriptional pathways labeled by our RAM system (Sørensen et al., 2016). If this hypothesis is correct, it will yield a new perspective regarding the CTA-conditioning active ensemble, CTA-memory retrieval, and memory generalization.

## 5 Data Availability Statement

All data generated from these experiments are included in the figures and supplemental figures. Raw data supporting the conclusions of this article will be made available by the authors upon request. Custom MATLAB scripts used for the analyses of electrophysiology data can be found at https://github.com/BrianAndCary/papers/tree/master/bcary2020_paper/mini_FI_GUI.

## 6 Author Contributions

These experiments were conceptualized by RR, CHW, and GGT. The behavior experiments were carried out by RR and CHW. RR performed the electrophysiology experiments, immunofluorescence staining, and analysis. RR wrote the original draft of this manuscript. RR, CHW, and GGT reviewed and edited the manuscript together.

## 7 Funding

This work was funded by NIH grant R35NS 111562 (GGT) and NS108506 (RR).

## 8 Conflict of Interest

The authors declare that the research was conducted in the absence of any commercial or financial relationships that could be construed as a potential conflict of interest.

## 9 Acknowledgments

We thank Don Katz, Bradly Stone, Jian-You Lin and members of the Turrigiano Lab for critical feedback on these experiments. Additionally, we thank Arianna Maffei for constructive feedback on an earlier draft of this manuscript.

## Notes

### Competing Interest Statement

The authors have declared no competing interest.

### Summary of Updates

This revision is meant to fix an error with the previous version of the manuscript in which an incorrect version of Figure 1 was uploaded.

## References

Allen, G. V., Saper, C. B., Hurley, K. M., & Cechetto, D. F. (1991). Organization of visceral and limbic connections in the insular cortex of the rat. The Journal of Comparative Neurology, 311(1), 1–16.

Angulo, R. (2018). Pre-exposure Schedule Effects on Generalization of Taste Aversion and Palatability for Thirsty and Not Thirsty Rats. Frontiers in Psychology, 9, 878.

Arieli, E., Younis, N., & Moran, A. (2021). Distinct progressions of neuronal activity changes underlie the formation and consolidation of a gustatory associative memory. The Journal of Neuroscience: The Official Journal of the Society for Neuroscience. https://doi.org/10.1523/JNEUROSCI.1599-21.2021

Asok, A., Kandel, E. R., & Rayman, J. B. (2019). The Neurobiology of Fear Generalization. Frontiers in Behavioral Neuroscience, 12, 329.

Avenet, P., & Lindemann, B. (1988). Amiloride-blockable sodium currents in isolated taste receptor cells. In The Journal of Membrane Biology (Vol. 105, Issue 3, pp. 245–255). https://doi.org/10.1007/bf01871001

Baird, J.-P., St John, S. J., & Nguyen, E. A.-N. (2005). Temporal and qualitative dynamics of conditioned taste aversion processing: combined generalization testing and licking microstructure analysis. Behavioral Neuroscience, 119(4), 983–1003.

Bures, J., Bermúdez-Rattoni, F., & Yamamoto, T. (1998). Conditioned taste aversion: Memory of a special kind. Oxford Psychology Series No. 31., 178.

Cary, B. A., & Turrigiano, G. G. (2021). Stability of neocortical synapses across sleep and wake states during the critical period in rats. eLife, 10. https://doi.org/10.7554/eLife.66304

Chotro, M. G., & Alonso, G. (1999). Effects of stimulus preexposure on the generalization of conditioned taste aversions in infant rats. Developmental Psychobiology, 35(4), 304–317.

Domjan, M. (1975). Poison-induced neophobia in rats: Role of stimulus generalization of conditioned taste aversions. Animal Learning & Behavior, 3(3), 205–211.

Dunsmoor, J. E., & Paz, R. (2015). Fear Generalization and Anxiety: Behavioral and Neural Mechanisms. Biological Psychiatry, 78(5), 336–343.

Flores, V. L., Parmet, T., Mukherjee, N., Nelson, S., Katz, D. B., & Levitan, D. (2018). The role of the gustatory cortex in incidental experience-evoked enhancement of later taste learning. Learning & Memory, 25(11), 587–600.

Frank, M. E., & Nowlis, G. H. (1989). Learned aversions and taste qualities in hamsters. Chemical Senses, 14(3), 379–394.

Gallo, M., Roldan, G., & Bures, J. (1992). Differential involvement of gustatory insular cortex and amygdala in the acquisition and retrieval of conditioned taste aversion in rats. Behavioural Brain Research, 52(1), 91–97.

Haley, M., Bruno, S., Fontanini, A., & Maffei, A. (2020). LTD at amygdalocortical synapses as a novel mechanism for hedonic learning. eLife.

Haley, M. S., Fontanini, A., & Maffei, A. (2016). Laminar- and Target-Specific Amygdalar Inputs in Rat Primary Gustatory Cortex. The Journal of Neuroscience: The Official Journal of the Society for Neuroscience, 36(9), 2623–2637.

Heck, G. L., Mierson, S., & DeSimone, J. A. (1984). Salt taste transduction occurs through an amiloride-sensitive sodium transport pathway. Science, 223(4634), 403–405.

Hengen, K. B., Torrado Pacheco, A., McGregor, J. N., Van Hooser, S. D., & Turrigiano, G. G. (2016). Neuronal Firing Rate Homeostasis Is Inhibited by Sleep and Promoted by Wake. Cell, 165(1), 180–191.

Heyer, B. R., Taylor-Burds, C. C., Tran, L. H., & Delay, E. R. (2003). Monosodium glutamate and sweet taste: generalization of conditioned taste aversion between glutamate and sweet stimuli in rats. Chemical Senses, 28(7), 631–641.

Josselyn, S. A., & Tonegawa, S. (2020). Memory engrams: Recalling the past and imagining the future. Science, 367(6473).

Kandel, E. R. (2012). The molecular biology of memory: cAMP, PKA, CRE, CREB-1, CREB-2, and CPEB. Molecular Brain, 5, 14.

Kandel, E. R., Dudai, Y., & Mayford, M. R. (2014). The molecular and systems biology of memory. Cell, 157(1), 163–186.

Kandel, E. R., Schwartz, J. H., Jessell, T. M., Siegelbaum, S. A., & Hudspeth, A. J. (2013). Principles of Neural Science, Fifth Edition. McGraw Hill Professional.

Kheirbek, M. A., Klemenhagen, K. C., Sahay, A., & Hen, R. (2012). Neurogenesis and generalization: a new approach to stratify and treat anxiety disorders. Nature Neuroscience, 15(12), 1613–1620.

Kiefer, S. W., & Braun, J. J. (1977). Absence of differential associative responses to novel and familiar taste stimuli in rats lacking gustatory neocortex. Journal of Comparative and Physiological Psychology, 91(3), 498–507.

Kiefer, S. W., & Braun, J. J. (1979). Acquisition of taste avoidance habits in rats lacking gustatory neocortex. Physiological Psychology, 7(3), 245–250.

Li, W.-G., Liu, M.-G., Deng, S., Liu, Y.-M., Shang, L., Ding, J., Hsu, T.-T., Jiang, Q., Li, Y., Li, F., Zhu, M. X., & Xu, T.-L. (2016). ASIC1a regulates insular long-term depression and is required for the extinction of conditioned taste aversion. Nature Communications, 7, 13770.

Lin, J.-Y., Roman, C., Arthurs, J., & Reilly, S. (2012). Taste neophobia and c-Fos expression in the rat brain. Brain Research, 1448, 82–88.

Maffei, A., Haley, M., & Fontanini, A. (2012). Neural processing of gustatory information in insular circuits. Current Opinion in Neurobiology, 22(4), 709–716.

Mahan, A. L., & Ressler, K. J. (2012). Fear conditioning, synaptic plasticity and the amygdala: implications for posttraumatic stress disorder. Trends in Neurosciences, 35(1), 24–35.

McLaren, I. P. L., & Mackintosh, N. J. (2002). Associative learning and elemental representation: II. Generalization and discrimination. Animal Learning & Behavior, 30(3), 177–200.

Miska, N. J., Richter, L. M., Cary, B. A., Gjorgjieva, J., & Turrigiano, G. G. (2018). Sensory experience inversely regulates feedforward and feedback excitation-inhibition ratio in rodent visual cortex. eLife, 7. https://doi.org/10.7554/eLife.38846

Moran, A., & Katz, D. B. (2014). Sensory cortical population dynamics uniquely track behavior across learning and extinction. The Journal of Neuroscience: The Official Journal of the Society for Neuroscience, 34(4), 1248–1257.

Mukherjee, N., Wachutka, J., & Katz, D. B. (2019). Impact of precisely-timed inhibition of gustatory cortex on taste behavior depends on single-trial ensemble dynamics. eLife, 8.

Nachman, M., & Ashe, J. H. (1973). Learned taste aversions in rats as a function of dosage, concentration, and route of administration of LiCl. Physiology & Behavior, 10(1), 73–78.

Nakashima, M., Uemura, M., Yasui, K., Ozaki, H. S., Tabata, S., & Taen, A. (2000). An anterograde and retrograde tract-tracing study on the projections from the thalamic gustatory area in the rat: distribution of neurons projecting to the insular cortex and amygdaloid complex. In Neuroscience Research (Vol. 36, Issue 4, pp. 297–309). https://doi.org/10.1016/s0168-0102(99)00129-7

Parker, L. A., & Revusky, S. (1982). Generalized conditioned flavor aversions: Effects of toxicosis training with one flavor on the preference for different novel flavors. Animal Learning & Behavior, 10(4), 505–510.

Ramirez, M.L. (2020). UnivarScatter. https://github.com/manulera/UnivarScatter.

Reilly, S., & Schachtman, T. R. (2008). Conditioned Taste Aversion: Neural and Behavioral Processes. Oxford University Press.

Richardson, R., Williams, C., & Riccio, D. C. (1984). Stimulus generalization of conditioned taste aversion in rats. Behavioral and Neural Biology, 41(1), 41–53.

Shepard, R. N. (1987). Toward a universal law of generalization for psychological science. Science, 237(4820), 1317–1323.

Smith, D. V., & Theodore, R. M. (1984). Conditioned taste aversions: generalization to taste mixtures. Physiology & Behavior, 32(6), 983–989.

Smith, K. R., Treesukosol, Y., Paedae, A. B., Contreras, R. J., & Spector, A. C. (2012). Contribution of the TRPV1 channel to salt taste quality in mice as assessed by conditioned taste aversion generalization and chorda tympani nerve responses. American Journal of Physiology. Regulatory, Integrative and Comparative Physiology, 303(11), R1195–R1205.

Sørensen, A. T., Cooper, Y. A., Baratta, M. V., Weng, F.-J., Zhang, Y., Ramamoorthi, K., Fropf, R., LaVerriere, E., Xue, J., Young, A., Schneider, C., Gøtzsche, C. R., Hemberg, M., Yin, J. C., Maier, S. F., & Lin, Y. (2016). A robust activity marking system for exploring active neuronal ensembles. eLife, 5. https://doi.org/10.7554/eLife.13918

Sun, X., Bernstein, M. J., Meng, M., Rao, S., Sørensen, A. T., Yao, L., Zhang, X., Anikeeva, P. O., & Lin, Y. (2020). Functionally Distinct Neuronal Ensembles within the Memory Engram. Cell, 181(2), 410-423.e17.

Takeuchi, T., Duszkiewicz, A. J., & Morris, R. G. M. (2014). The synaptic plasticity and memory hypothesis: encoding, storage and persistence. In Philosophical Transactions of the Royal Society B: Biological Sciences (Vol. 369, Issue 1633, p. 20130288). https://doi.org/10.1098/rstb.2013.0288

Ting, J. T., Daigle, T. L., Chen, Q., & Feng, G. (2014). Acute Brain Slice Methods for Adult and Aging Animals: Application of Targeted Patch Clamp Analysis and Optogenetics. In M. Martina & S. Taverna (Eds.), Patch-Clamp Methods and Protocols (pp. 221–242). Springer New York.

Torrado Pacheco, A., Bottorff, J., Gao, Y., & Turrigiano, G. G. (2021). Sleep Promotes Downward Firing Rate Homeostasis. Neuron, 109(3), 530-544.e6.

Wu, C.-H., Ramos, R., Katz, D. B., & Turrigiano, G. G. (2021). Homeostatic synaptic scaling establishes the specificity of an associative memory. Current Biology: CB, 31(11), 2274-2285.e5.

